# IG-28 enables identification and quantification of high-confidence isoforms from large bulk and single-cell full-length cDNA datasets

**DOI:** 10.64898/2026.10.23.761370

**Authors:** Alessandro Calicchio, Amanda O. Li, Tanvi Herwadkar, Honey E. Mekonen, Christopher Vollmers

## Abstract

The technology to sequence full-length transcriptomes from bulk and single-cell samples has rapidly advanced. As a result, full-length cDNA datasets now reach billions of reads in size which is beyond the capability of most established isoform identification tools. Here, we introduce IG-28 which addresses this challenge with its capability to process and analyze large fl-cDNA datasets fast and using minimal computational resources.

In contrast to its predecessor Mandalorion, IG-28 is not limited to bulk fl-cDNA but is capable of parsing, processing, and analyzing single-cell fl-cDNA datasets. It does so in a lightweight, flexible manner, integrating seamlessly with established scRNA-seq analysis workflows.

We benchmarked IG-28 against a variety of established and new isoform identification tools using several bulk and a complex single-cell fl-cDNA dataset and found that IG-28 consistently outperformed the other tools by identifying high-confidence ‘public’ isoforms.

## INTRODUCTION

The technology for capturing RNA transcripts end-to-end in the form of full length (fl) cDNA sequencing has improved dramatically in the last decade^1,2^. In particular, the PacBio Kinnex^3^ and ONT-based R2C2^4,5^ methods now allow for the sequencing of tens to hundreds of millions of highly accurate fl-cDNA reads in single sequencing runs. However, conclusively identifying transcript isoforms based on the fl-cDNA reads these methods produce remains a challenge.

While many tools exist for this purpose^6–11^ and have been benchmarked by their developers, consortia like LRGASP have shown that most tools identify very different sets of transcript isoforms when analyzing the same fl-cDNA reads^12^. Considering fl-cDNA reads cover entire transcripts – all splice junctions as well as transcription start sites (TSS) and polyA sites – the fact that tools for isoform identification from fl-cDNA reads generally don’t agree on the sets of transcript isoforms they identify is a major concern as it undermines the confidence in each tool’s output.

Further, most tools independently benchmarked by LRGASP weren’t designed for and therefore often can’t handle fl-cDNA datasets containing hundreds of millions or even billions of fl-cDNA reads. However, the increase in throughput, in particular by the PacBio Kinnex method, now means that datasets of this size are becoming the standard rather than a rare exception. While previous efforts have circumvented this limitation of tools by breaking datasets up by chromosomes^13^, this is at best a temporary solution until even individual chromosomes will be covered by hundreds of millions of fl-cDNA reads.

Finally, only a few tools currently available are capable of parsing and analyzing fl-cDNA data generated using single-cell transcriptomics protocols^8,14,15^. However, fl-cDNA based analysis of single-cells transcriptomics experiments is becoming more common because most high-throughput single-cell RNA-seq methods, including 10X Genomics and Parse, generate fl-cDNA as an intermediate product of their short-read RNAseq workflow. This fl-cDNA intermediate product can be analyzed by the increasingly powerful long-read sequencing methods.

Here we introduce the IG-28 tool, the successor of our Mandalorion tool^16^. IG-28 retains some parts of Mandalorion’s isoform identification logic but features heavily reworked code base. As a result, IG-28 rapidly identifies and quantifies truly high-confidence transcript isoforms from fl-cDNA read datasets billions of reads in size while requiring minimal computing resources. Further, IG-28 can parse single-cell fl-cDNA data to create pseudo-bulk transcriptomes of individual cell types.

To benchmark IG-28 we analyzed bulk fl-cDNA data from 1) an Iso-seq dataset covering the transcriptome of WTC-11 cell-line produced for the LRGASP effort featuring SIRV spike-ins, 2) a combination of a public PacBio Kinnex dataset and an ONT-based R2C2 dataset produced for this study covering the human brain transcriptome 3) a public PacBio Kinnex dataset covering 100 lymphoblastoid cell lines of the 1000 Genomes Project. We also established IG-28’s ability to parse and analyze single-cell fl-cDNA datasets by pooling and processing multiple public PacBio Kinnex fl-cDNA datasets containing ∼100,000 human peripheral blood mononuclear cells (PBMCs).

We then compared the isoform sets IG-28 identified to ground truths and to the isoform sets called by other tools. For this purpose we defined a new metric called *% public SJCs* that treats splice-junction chains identified by more than one tool as high-confidence. This comparison established IG-28 as a fast and memory efficient tool identifying and quantifying high-confidence isoforms in both bulk and single-cell fl-cDNA sequencing data.

## RESULTS

### IG-28 bulk fl-cDNA workflow

As the successor to Mandalorion, IG-28 retains some of the logic that Mandalorion used to filter isoforms. As such, we expect isoforms reported by IG-28 to have a strong overlap with those reported by Mandalorion on the same dataset. However, to reduce runtime and RAM usage as well as enable the analysis of single cell fl-cDNA datasets, IG-28 represents a major rewrite from Mandalorion for how isoforms are identified and quantified.

For bulk fl-cDNA analysis, IG-28 is separated into modules that align the reads (module A), parse reads to identify, filter, and quantify isoforms (module D), and add CDSs to each isoform (module C) (Fig. 1). By default, IG-28 runs modules A, D, and C sequentially. Module A accepts a sample sheet where each line represents an input file location (fasta/q or bam files) and its associated sample and group names in separate columns. IG-28 combines the reads contained in all input files into a single fasta file while modifying each read name with sample and group name like this:

{SampleName}|{GroupName}|{OriginalReadName}

**Fig. 1:**
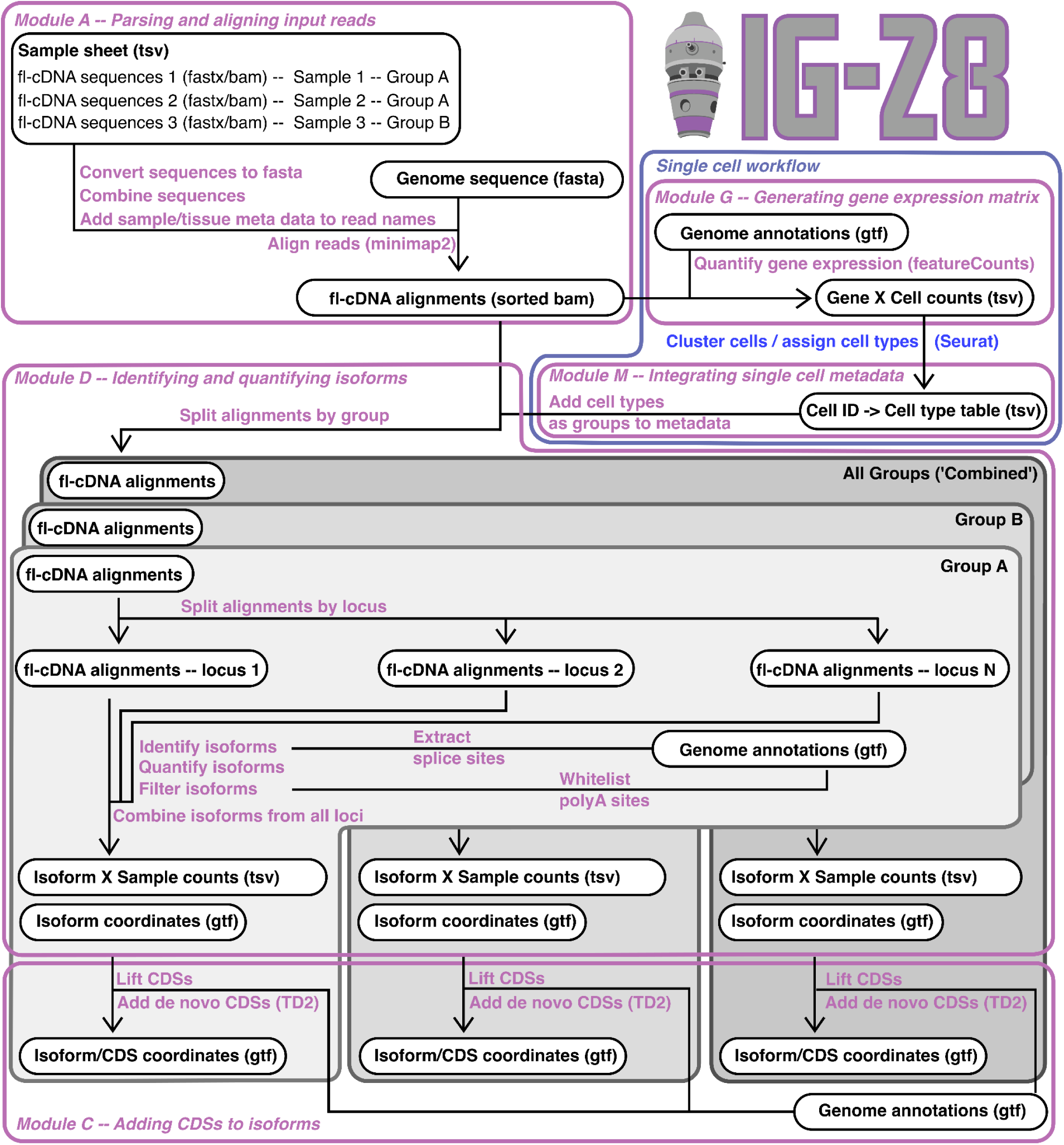
IG-28 workflow. IG-28 analyses bulk and single-cell fl-cDNA data. Its workflow is divided into modules. Module A parses input data and aligns reads to the reference genome using minimap2. Module D, splits alignments by group, if information about them is provided, then processes all and the group-separated alignments sequentially. It does so by splitting reads by locus, then identifies splice sites, links them into SJCs, that are then assigned TSS and polyA sites and filtered to generate and quantify isoforms. Modules G and M, can be used during single-cell data processing. Module G produces a gene expression data matrix for single-cell clustering by tools like Seurat or Scanpy, while module M imports the resulting cell-to cell type assignment back in IG-28. Module C adds CDSs to identified isoforms.

By retaining sample and group information in this way, module D of IG-28 can – quickly and using minimal RAM as well as read/write operations – identify isoforms on the group level and then quantify these isoforms on the sample level. IG-28 does so by first splitting all reads by group and then processing each group individually and sequentially. For each group, it splits all the reads for that group into non-overlapping loci. It then processes loci for each group in a multithreaded manner (Fig. 1). IG-28 identifies and quantifies isoforms by first grouping reads by their splice junction chain (SJC). For each SJC, IG-28 then defines TSS and polyA site locations based on the ends of the reads assigned to the SJC. Those reads are then split into TSS-SJC-polyA groups, i.e. isoforms. These isoforms are filtered based on their length, read number, and whether or not they are likely to be an artifact.

In addition to group level outputs, IG-28 also creates a ‘Combined’ output by combining all groups present in the dataset.

At this point, IG-28 reports its results in subdirectories named after each group (as well one named ‘Combined’). Within each subdirectory it generates:

1) a gtf file containing the structure of all identified isoforms as well as
2) a tab-delimited text file containing the read counts for each isoform in each sample.

To evaluate protein-coding potential of the isoforms IG-28 identifies, module C then lifts open reading frames (ORFs) from the reference annotation (if provided) and uses the external TD2^17^ tool to call ORFs *de-novo* for each isoform. It then adds the longest ORF (either lifted or called *de-novo*) to each isoform as part of a modified, additional gtf file (Fig. 1).

### Characteristics of bulk full-length cDNA datasets used for comparisons

To test its isoform identification and quantification performance, we applied IG-28 to bulk fl-cDNA datasets of varying sizes (Table 1).

**Table 1:** Bulk fl-cDNA dataset characteristics.

| Data set | Sequencing Method | Read numbers |
| --- | --- | --- |
| **WTC-11 & SIRV** | Iso-seq\|PacBio | 7,424,923 |
| **Human Brain** | Kinnex\|PacBio | 20,138,817 |
|  | R2C2\|ONT | 8,633,082 |
| **1KG** | Kinnex\|PacBio | 1,005,284,663 |

The first dataset (**WTC-11 & SIRV**) contained ∼7 million PacBio Iso-seq reads covering the transcriptome of the human WTC-11 cell line. This dataset, which was generated for the LRGASP consortium and is hosted by ENCODE^18^, also contained reads covering SIRV set 4 spike-in RNA molecules with a known ground-truth.

The second bulk dataset (**Human Brain**) contained a total of about ∼30 million PacBio Kinnex and R2C2 (ONT) reads covering the transcriptome of the human brain. While the PacBio Kinnex reads in this dataset were generated by PacBio, the R2C2 reads were generated by us specifically for this study using a size-selection strategy resulting in samples with distinct read-length distributions.

The third bulk dataset (**1KG**) contained about ∼1 billion PacBio Kinnex reads covering the transcriptomes of a subset of the cell lines used in the 1000 Genome Project^19^. This dataset is part of an ongoing 1000 Genomes long-read RNA-seq project and the full dataset is unpublished and in preparation (1000 Genomes Long-Read Sequencing Consortium).

Together, these 4 datasets represent a wide range of use cases by featuring different read depths, read lengths, and sample numbers.

### IG-28 accurately detects spike-in isoforms

To first evaluate IG-28 performance on a dataset with a known ground-truth, we analyzed the *WTC-11 & SIRV* dataset of standard Iso-seq PacBio fl-cDNA sequencing reads. This data set has exceptional quality, features a wide read length distribution and has been analyzed by many tools as part of LRGASP and beyond. Importantly, it also contains SIRV set 4 spike-in transcripts.

In addition to IG-28 (v1.0), we also processed the *WTC-11 & SIRV* data set using the long established and independently benchmarked tools *Mandalorion* (v4.6), *FLAIR (v3.0.0)*^20^*, and IsoQuant (v3.12.0)* ^21^, as well as the recently released tools *LRAA (v0.15.0)*^8^, and the PacBio-developed tool *Isocall (v0.15.0)*^22^.

We ran all tools using default settings (for HiFi reads if appropriate) using the GRCh38 primary assembly human reference sequence. We ran all tools with the GENCODE v49 human reference annotation plus the annotation for the SIRV set 4 spike-ins. Further, we also ran the tools that allowed it (IG-28, Mandalorion, IsoQuant, and LRAA) without providing GENCODE v49 reference annotation.

First, we evaluated the performance of all tools when identifying SIRV isoforms (Fig. 2A). When provided with an annotation – which for the SIRV transcripts represents the ground truth – all tools performed very well. FLAIR in particular detected all SIRV True Positives with only one False Positive. IG-28 performed slightly better than its predecessor Mandalorion. IG-28 detected 64 out of 69 SIRVs with the ones being missed mostly being very short single exon isoforms that are outside of IG-28 default isoform length cut-offs. The only False Positive IG-28 detected was an isoform with a very short terminal exon that can’t be aligned by minimap2 and is therefore omitted in the isoform identified by IG-28. LRAA and Isocall also performed well in identifying True Positive SIRV isoforms. However, both tools showed a comparatively high number of False Positives. In the case of LRAA, that was exaggerated in the absence of a reference annotation. IsoQuant, while detecting no False Positives with annotation, showed the largest number of False Positives when run without annotation.

**Fig. 2:**
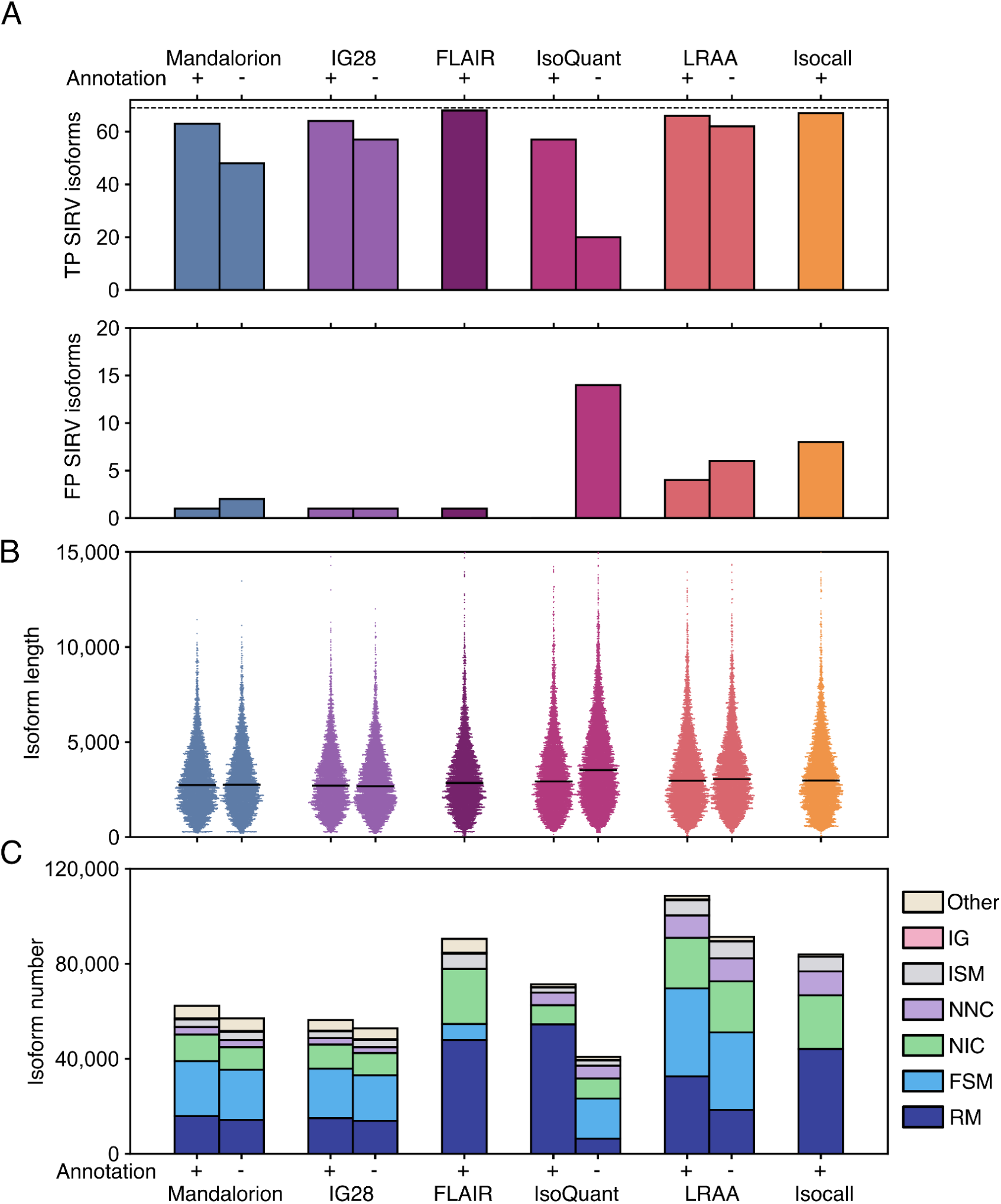
Comparison of identified isoforms in the *WTC-11 & SIRV* dataset. A). Upper panel: number of true positive (TP) SIRV transcripts identified by each tool. Lower panel: number of false positives (FP) SIRV transcripts identified by each tool. B). Comparison of length distribution of the isoforms detected by each tool. The black line shows the median read length. C). Classifications of isoforms generated by each tool. Where possible, the representations show isoform identified with and without providing a reference annotation.

In summary, IG-28 performed very well when detecting SIRV isoforms, balancing Precision and Recall without over-reliance on a reference annotation.

### IG-28 output is stable when run without a provided reference annotation

Moving beyond spike-in transcripts we compared the isoforms different tools identified for the entire *WTC-11 & SIRV* data set. Here, with the exception of IsoQuant run without an annotation, all tools called isoforms of overall similar length (Fig. 2B). Isoform numbers varied between ∼40,000 (IsoQuant without annotation) to ∼110,000 (LRAA with annotation) (Fig. 2B).

Using SQANTI, we categorized these isoforms into the following categories: ‘full splice match’ (FSM, when the entire splice junction chain (SJC) of an identified isoform matches the SJC of a reference isoform), ‘novel in catalog’ (NIC, when an identified isoform contains only splice sites present in the reference annotation but those splice sites are combined into at least one splice junction that is not found in the reference annotation), ‘novel not in catalog’ (NNC, when the an identified isoform contains at least one splice sites that is not present in the reference annotation), ‘incomplete splice match’ (ISM, when the SJC of an identified isoform is a partial match to the SJC of a reference isoform), intergenic (IG, isoforms found in intergenic regions), and Other (for all the transcripts that can’t be placed in one of the previous categories) (Fig. 2C). We also used a FSM subcategory called ‘reference match’ (RM) which represents FSM isoforms whose 5’ and 3’ ends closely match the reference isoform they are assigned to. This category is useful to identify TSS and polyA novelty within the FSM category. It also highlights tools which rely on reference annotations to determine the ends of their isoforms – in this case IsoQuant and Isocall.

IG-28 and Mandalorion isoform numbers and categories were similar whether an annotation was provided or not. On the other hand, IsoQuant and LRAA called considerably different numbers and categories of isoforms when not provided with an annotation (Fig. 2C).

For further analysis, we faced the challenge that the GENCODE v49 reference annotation does not represent a ground truth for isoforms actually present in *WTC-11 & SIRV*, *Human Brain*, and *1KG* datasets. Therefore, while we could determine a few descriptive metrics for each tool, we could not establish which one was more likely to detect correct, high-confidence isoforms. While simulated data could partially address this, simulation comes with its own issues. First, simulated data is still rooted in reference annotations which don’t reflect the actual transcriptome of any single tissue, cell type, or cell. Second, while the error profile of a sequencing technology can be modeled, modeling the actual molecular biology features and artifacts of fl-cDNA generation is a formidable challenge.

### IG-28 identifies public, high-confidence isoforms

To address the absence of a ground truth when evaluating isoform sets called by tools on different fl-cDNA datasets, we made one assumption: Identifying isoforms based on fl-cDNA reads is not an intrinsically hard problem because every fl-cDNA read presumably contains full information for the isoform it is derived from.

From this assumption follows that high-confidence isoforms should be called by more than one analysis tool (public). Second, and following from the first assumption, isoforms called by a single tool (private) are likely low-confidence isoforms.

To establish which tools in our study detected the most public, high-confidence isoforms, we compared the splice junction chains detected by each tool. We chose SJCs because IsoQuant and Isocall relied on the GENCODE reference annotation to define isoform ends which meant they no longer represented the experimental data.

For each combination of tools, we calculated what percentage of SJCs called by Tool A were also called by Tool B and what percentage of SJCs called by Tool B were also called by Tool A. In this way, we hoped to provide a clearer picture of how the outputs of different tools are related.

This analysis showed that, as expected, SJCs called by IG-28 were largely called by Mandalorion as well and vice versa. Further 92% and 95% of SJCs called by IG-28 were also called by LRAA and Isocall, respectively. However, only about 60% of SJCs called by LRAA were called by Isocall and vice versa. SJCs of FLAIR and IsoQuant were somewhat similar to each other, with 41% of SJCs called by FLAIR SJCs being also called by IsoQuant SJCs and 45% of SJCs called by IsoQuant also being called by FLAIR.

Next, we calculated what percentage of SJCs called by each tool was also called by at least one other tool, i.e. what percentage of their SJCs were public. Here, we didn’t include IG-28 and Mandalorion in each other’s calculation as their outputs are related and therefore highly similar. We found that 99% of SJCs called by IG-28 were public. This was followed by Mandalorion (96%), Isocall (75%), LRAA (68%), FLAIR (62%), and Isoquant (55%) (Fig. 3A, left panel).

**Fig. 3:**
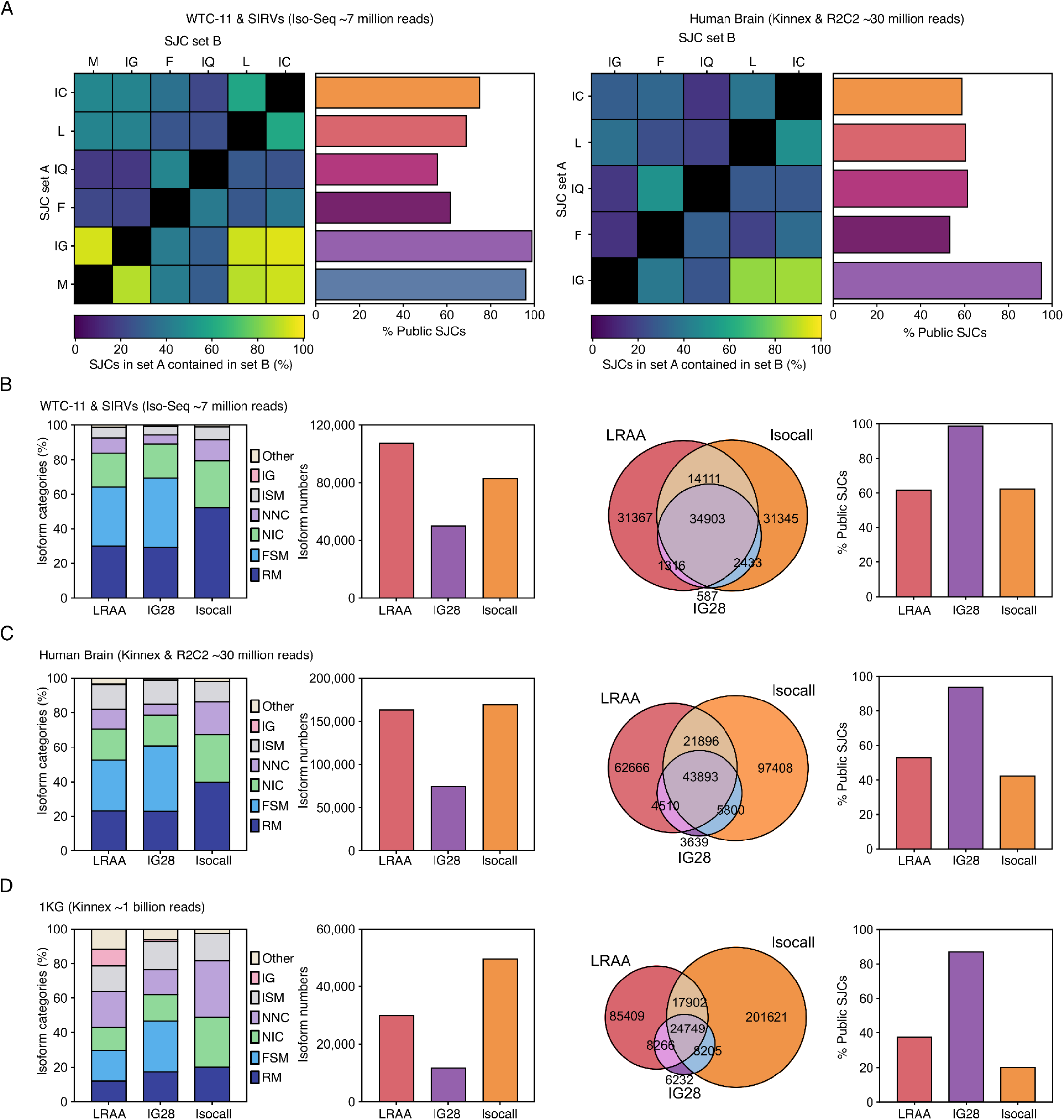
IG-28 identifies public ‘high confidence’ isoforms. A) The heatmaps show the pairwise comparison between the splice junction chain (SJC) identified by tools. A brighter color indicates a higher percentage of SJC identified by tool A are also described by tool B. The barplots show the percentage of SJC identified by one tool that is also identified by at least one other tool, i.e. public. Data for the *WTC-11 & SIRV* dataset are on the left, data for the *Human Brain* dataset are on the right. B-D) Comparison between the transcriptomes generated by LRAA, IG-28 and Isocall for the *WTC-11 & SIRV* (B), *Human Brain* (C) and *1KG* (D) datasets. Panels show, from left to right, isoform categories, the number of isoforms, the overlap between their SJC sets and the percentage of public SJCs (SJCs identified by at least another of the three tools).

To investigate whether this behavior was unique to the *WTC11 & SIRV* data set, we next called isoforms for the 30 million fl-cDNA read *Human Brain* dataset composed of both PacBio Kinnex and size-selected R2C2 reads (Fig. 3A, right panel). Comparing the SJCs called by the different tools (we did not include Mandalorion in further analysis) in the same way, we found that again, SJCs called by IG-28 were also called at very high rates by Isocall and LRAA. Further, 95% of IG-28 isoforms were public while other tools showed a percentage of around 60% public SJCs.

Not only were the SJCs called by IG-28 also often called by LRAA and Isocall, SJCs called by IG-28 tended to be called by both LRAA and Isocall (Fig. 3B). To rephrase this: IG-28 called the SJCs that LRAA and Isocall agreed on.

This relationship held up across *WTC11 & SIRV*, *Human Brain*, and *1KG* data sets (for the *1KG* dataset, comparison was limited to chr1 as LRAA would have taken several weeks to process the entire datasets on our workstations). As the size of datasets increased, the relative overlap of LRAA and Isocall decreased. However, 97%, 92%, and 85% SJCs called by IG-28 fell within the SJCs called by either LRAA or Isocall in the *WTC11 & SIRV*, *Human Brain*, and *1KG* datasets respectively (Fig. 3B, 3C, and 3D). Finally, as intergenic and NNC isoforms increase strongly in LRAA and Isocall output of the 1KG dataset, respectively, isoform categories produced by IG-28 changed far less compared to smaller datasets.

### IG-28 quantification is concordant with Isocall quantification

All the tools we compare in this study also quantify the isoforms they identify. To benchmark the ability of these tools to quantify isoforms, we chose to perform pairwise comparison of the counts assigned to each SJCs called by both tools (Fig. 4A).

**Fig. 4:**
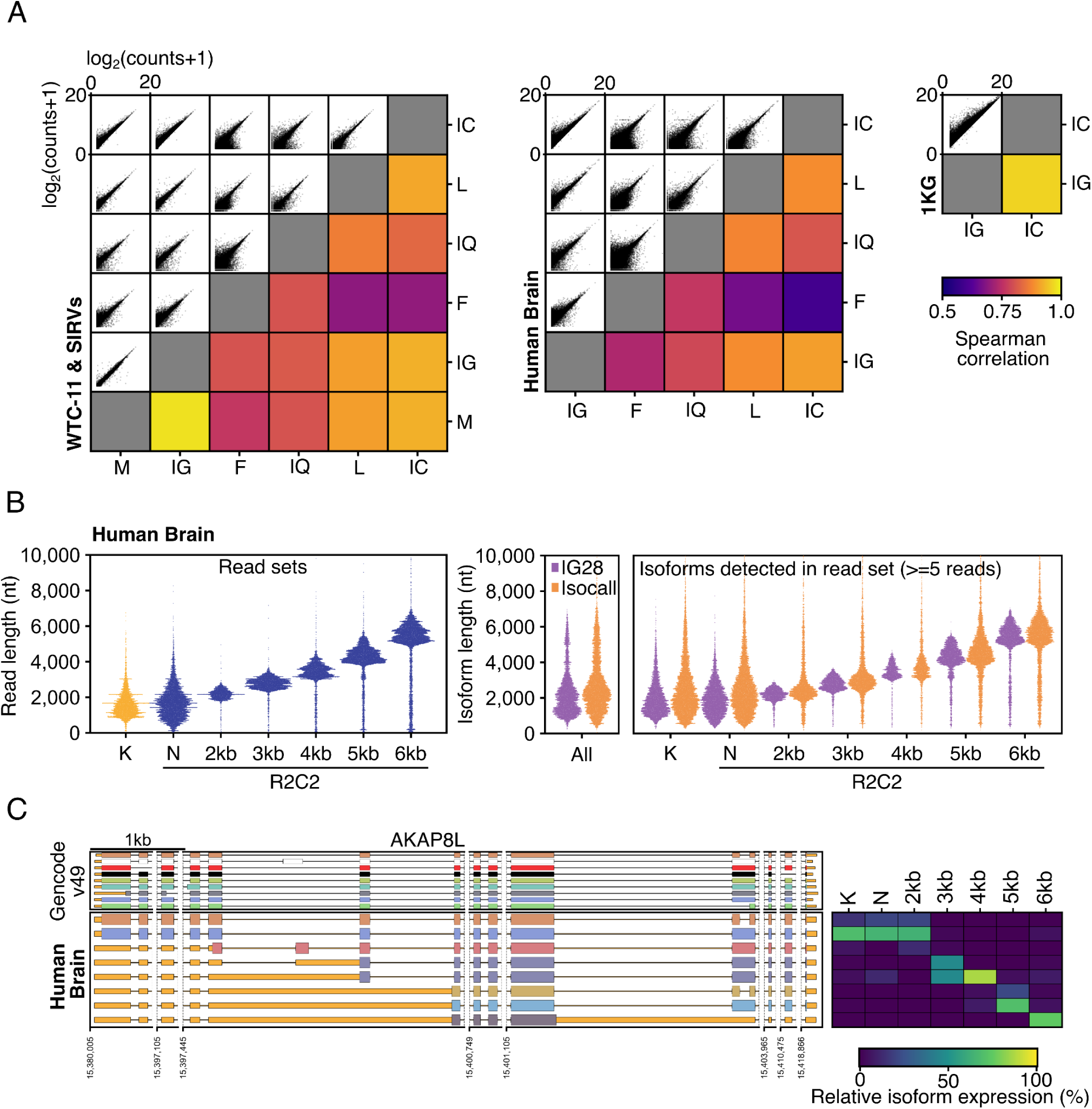
IG-28 quantifies isoform at the sample level. A) Pairwise comparison of read counts assigned to each SJC by different tools. From left to right, data from the *WTC-11 & SIRV, Human Brain*, and 1KG dataset are shown. (M: Mandalorion; IG: IG-28; F: FLAIR; IQ: IsoQuant; L: LRAA; IC: Isocall) B) The left panel shows read length distribution in the Human Brain sample. This dataset contains reads obtained from different library preparation protocols: Kinnex (K, in yellow), and R2C2 (in blue). R2C2 was used to sequence the same cDNA library before (non-size selected or N) and after size selection of fl-cDNA molecules (5 size bins processed: 2kb, 3kb, 4kb, 5 kb and 6kb). The center panel shows the isoform length distribution for isoforms identified by IG-28 and Isocall. The right panel shows which of the isoforms in the center panel contained at least 5 reads from the indicated Kinnex and R2C2 libraries. C) Genome browser plot of AKAP8L gene locus showing both Gencode annotated isoforms (upper panel) and IG-28 identified isoforms (bottom panel) in the Human Brain dataset. The heatmap shows the relative expression of the 8 IG-28 defined isoforms in each 7 fl cDNA libraries.

Again, as expected, IG-28 showed high correlation with Mandalorion counts (Fig. 4A, left, center). Beyond that, IG-28 and Isocall showed the highest correlation for *WTC11 & SIRV* and *Human Brain* data sets. IG-28 and Isocall also showed very high correlation on the 1KG dataset, but were the only tools that completed that analysis making it impossible to compare to other tools (Fig. 4A left, center, right).

Interestingly, while IG-28 counts are highly correlated with Isocall, they are consistently lower. To investigate the cause of this, we focused on the *Human Brain* datasets which pairs publicly available PacBio Kinnex data and size-selected R2C2 data (Fig. 4B). The Kinnex and size-selected R2C2 data was provided to IG-28 and Isocall as independent samples. As such, both tools identified a single set of isoforms for the dataset but quantified these isoforms on the level of ‘samples’ – in this case Kinnex and size-selected R2C2 libraries.

This made it possible to connect the length of reads and the isoforms they are assigned to for quantification purposes. What we found was that IG-28 assigns reads to isoforms of the same length (Fig. 4B, right). This is due to a read having to match both all splice junctions and ends of an isoform to be counted towards it. A good example of this is the AKAP8L gene that through a combination of intron retention events expresses isoforms of highly variable length. As the length of an isoform increases, so does the relative expression of that isoform – as measured by IG28 – in the size-selected cDNA bins that match the length of that isoform (Fig. 4C).

Isocall on the other hand assigns reads to isoforms that are both longer or shorter than the reads themselves which suggests Isocall accounts for partially matching reads towards isoforms quantification (Fig. 4B, right). This could explain in part why Isocall shows higher read counts for shared SJCs.

Overall, this established that IG-28 quantifications are highly correlated with Mandalorion which we have previously shown to be accurate^16^ for isoform quantification. Further, of all pairwise tool combinations, IG-28 and Isocall showed the highest correlation, which, in the absence of a ground truth, might indicate convergence on the actual abundance of the identified isoforms. Finally, as visualized for the AKAP8L gne in the *Human Brain* dataset, IG-28 creates sample-level quantification of isoforms by default which form the basis for the downstream analysis, including detection of differential isoform expression.

### IG-28 single-cell fl-cDNA workflow

While we have used Mandalorion for single cell transcriptome analysis^5^, we did so by making extensive use of custom scripts making the workflow quite involved. To simplify single-cell transcriptome analysis with IG-28 we developed two additional modules – G and M – which are run after module A (read parsing and alignment) and before module D (group-level isoform identification).

The full workflow then works as follows:

For single-cell data, which is commonly stored in unaligned bam files, the sample sheet is formatted in the following way:

~~~
/path/to/sc-flcDNA1.bam scRNA;flcDNA1;CB None
~~~

Running Module A allows IG-28 to extract the cell ID from each read – in this case stored under the CB flag. This cell ID is then combined with the Experiment ID (e.g. “flcDNA1”) and used in place of a sample name in the read name of the combined fasta going into alignment – in essence, each single cell is treated as its own sample. After alignment, using module G, IG-28 uses the external tool featureCounts^23^ to assign each aligned read to a gene. IG-28 then uses cell ID information in each read name to generate a CellxGene expression matrix.

This matrix can then be used to cluster cells and generate a Cell-to-Cell Type metadata file using stand-alone tools like Seurat or scanpy. In this way, established short-read single-cell RNA-seq cell clustering and annotation workflows can easily be adapted to fl-cDNA approaches. IG-28 uses module M to parse this Cell-to-Cell Type metadata file to annotate read names (module M) in the aligned read bam file to adhere to this format:

~~~
{ExperimentID}_{CellID}|{CellType}|{OriginalReadName}
~~~

In this way, cells are used as samples and cell types. Module D will therefore identify isoforms on the level of cell types (pseudo-bulk) while quantifying each identified isoform at the level of cells.

As with bulk data, IG-28 does so by first splitting all reads by cell type and then processing each cell type individually and sequentially. For each cell type, it splits all the reads for that cell type into non-overlapping loci and identifies, filters, and quantifies isoforms for each locus in a fully multithreaded manner.

As with bulk data, in addition to cell type-level isoforms, IG-28 also creates a ‘Combined’ output, identifying isoforms based across all cell types and quantifying those isoforms across all cells in the dataset.

### IG-28 generates CellxGene matrices for complex single cell data

To test IG-28 single-cell analysis capabilities we applied it to the analysis of a combined single-cell PBMC dataset composed of 7 individual experiments performed by PacBio and provided freely at

~~~
https://downloads.pacbcloud.com/public/dataset/Kinnex-single-cell-RNA/
~~~

The individual experiments used different 10X Genomics chemistries^24^ processed different numbers of cell numbers, and were sequenced on different PacBio instruments (Table 2).

**Table 2.** Single-cell fl-cDNA datasets.

| Dataset name | 10X Genomics Chromium workflow | Cell numbers | Read number | Instrument |
| --- | --- | --- | --- | --- |
| 10x3p | Next GEM Single Cell 3' | 12,840 | 81,678,354 | Revio |
| 10x5p | Next GEM Single Cell 5' | 13,984 | 44,323,585 | Revio |
| 10kcells 10xGEMX3p | GEM-X Single Cell 3' v4 | 10,324 | 110,681,049 | Revio |
| 10kcells 10xGEMX3p Vega | GEM-X Single Cell 3' v4 | 10,329 | 54,980,296 | Vega |
| 20kcells 10xGEMX3p Rep1 | GEM-X Single Cell 3' v4 | 20,426 | 218,678,519 | Revio |
| 20kcells 10xGEMX3p Rep2 | GEM-X Single Cell 3' v4 | 20,218 | 225,778,491 | Revio |
| 20kcells 10xGEMX5p | GEM-X Single Cell 5' v3 | 17,654 | 197,544,312 | Revio |

Overall this single-cell dataset (**scPBMC**) contained about ∼900 million PacBio Kinnex reads covering the transcriptomes of more than 100,000 PBMCs.

The sample sheet structure and meta data handling of IG-28 allowed us to track fl-cDNA reads from different experiments. After running module A we confirmed that each dataset contained a variable number of (spliced) reads and read length distributions (Fig. 5A).

**Fig. 5:**
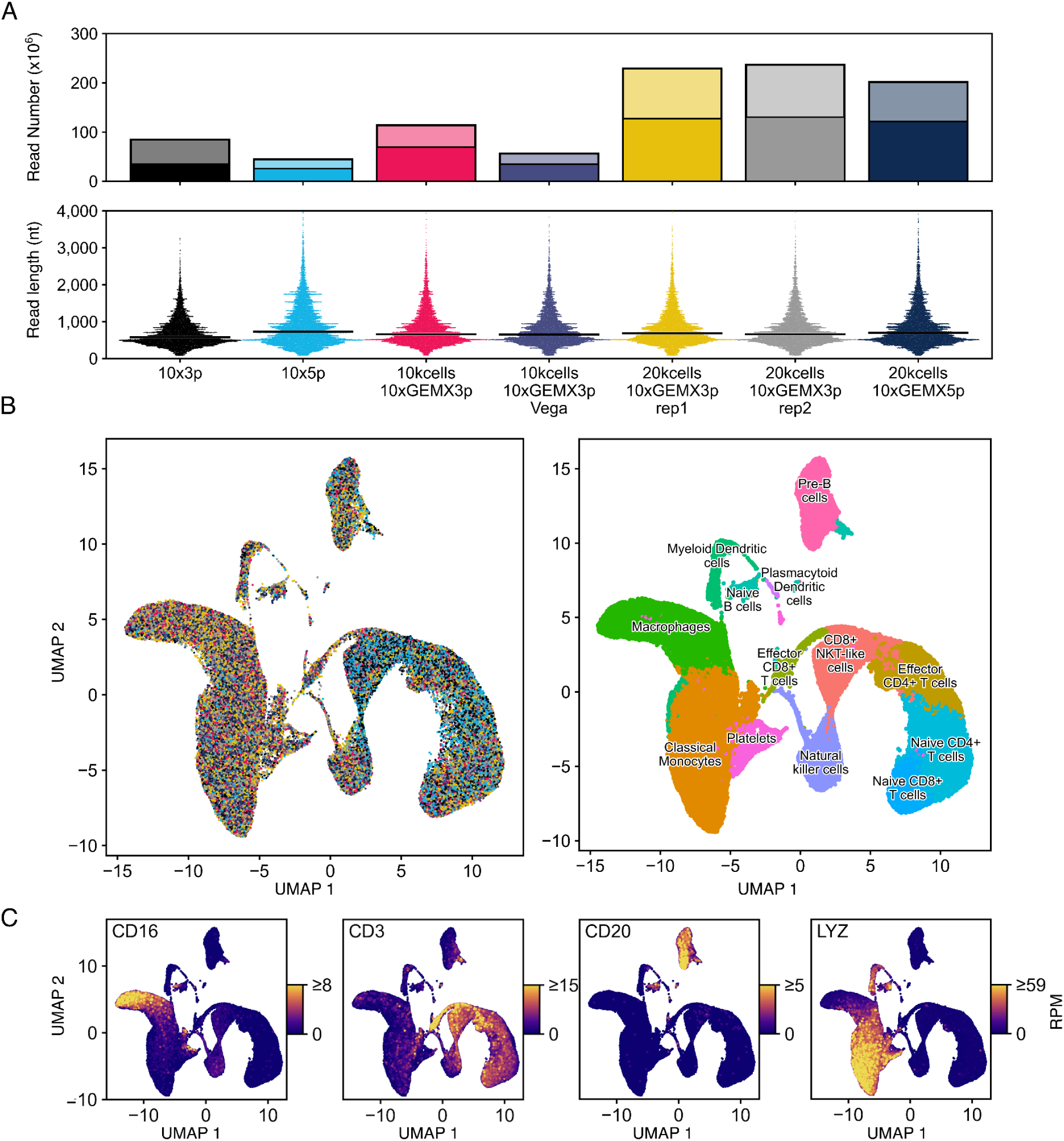
IG-28 combines and analyses multiple single-cell fl-cDNA data sets. A) The panels show the numbers and lengths of the aligned fl-cDNA reads in each of the 7 single-cell fl-cDNA Kinnex datasets. Numbers are shown as bar plots with faint bars indicating total aligned reads and solid bars indicating aligned reads with at least one splice junction. Read lengths are shown as swarm plots. B,C) All cells in the combined single cell data set are clustered using Seurat based on their gene expression profiles. The resulting UMAP visualizations show cells either colored by the individual dataset they originated from (B left, datasets colored as in A), the cell type they were annotated with (B right), or the expression of the indicated marker genes (C). For each gene, expression values were capped to the 99th percentile of expression across cells expressing that gene.

Running module G, IG-28 generated a CellxGene expression matrix for the combined experiments. We used this matrix as input into single cell RNA sequencing analysis with Seurat^25^. This expression matrix contained the experiment ID as part of each cell name, which allowed us to harmonize the combined dataset using batch correction with Canonical Correlation Analysis (CCA) integration^26^. After harmonization, we clustered cells and automatically assigned cell types with the scType method^27^.

To visualize the harmonized, clustered, and labeled dataset, we generated UMAPs, coloring cells either by experiment of origin (Fig. 5B, left panel) – observing the absence of experiment-specific clusters – or cell type assignment which showed 13 PBMC specific cell types (Fig. 5B, right panel). Importantly, the expression of CD16 (FCGR3A) CD3D, CD20 (MS4A1), and LYZ – known to be expressed by Macrophages, T cells, B cells, and Monocytes, respectively^28^ – was most pronounced in their associated cell types (Fig. 5C).

Overall, this established that the CellxGene matrix generated by IG-28 can be used even for complex single-cell analysis workflows.

### IG-28 generates pseudo-bulk transcriptomes from deep single cell fl cDNA data

Next, we exported a cell-to-cell type mapping file from the Seurat meta data. Using this cell-to-cell type file, we ran module M of IG-28 to modify experiment metadata and allow for the generation of celltype specific pseudo-bulk transcriptomes. Subsequently running modules D identified and quantified pseudo-bulk transcriptomes for each cell type.

As part of module D, IG-28 generates a bam file for each cell type in which read names carry information on their cell and experiment (run) of origin which makes quality control and downstream analysis straightforward. Parsing these bam files for each cell type showed that, as expected, cell types contained varying numbers of cells and (spliced) reads (Fig. 6A, top). Further parsing the data on the experiment level showed that different cell types had different read contributions from different datasets (Fig. 6A, bottom). The 10×3p and 10×5p experiments in particular were over-represented in T and Pre-B cells which is also visible in the UMAP with cells colored by experiment (Fig.6A, Fig. 5B left).

**Fig. 6:**
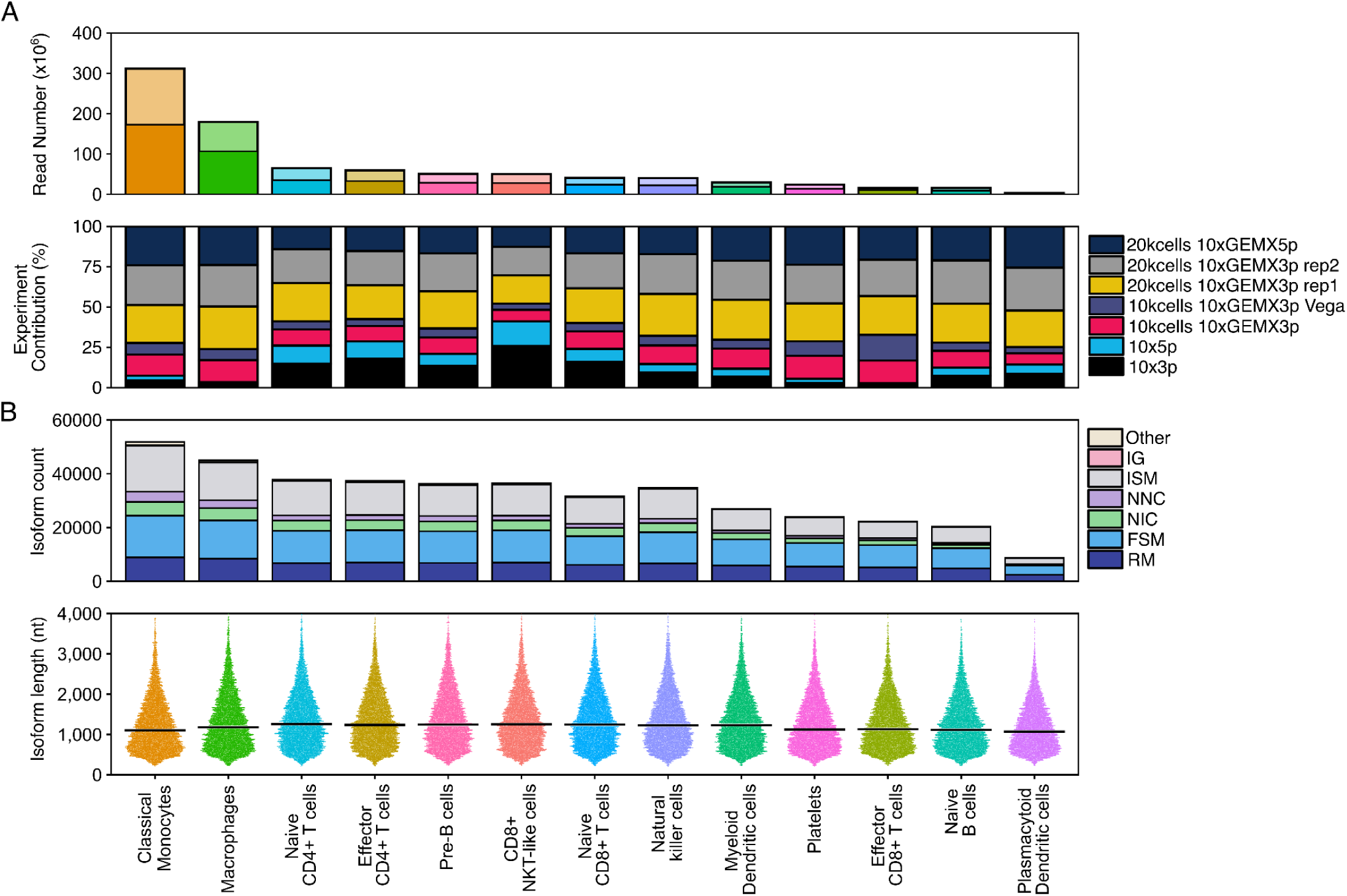
IG-28 generates cell type transcriptomes. A) The top panel shows the numbers of the aligned fl-cDNA reads for each annotated cell type. Numbers are shown as bar plots with faint bars indicating total aligned reads and solid bars indicating aligned reads with at least one splice junction. The bottom panel shows the relative contribution of the different single-cell fl-cDNA Kinnex datasets (runs) to the fl-cDNA reads of each cell type. B) The top panel shows the number of isoforms identified by IG-28 for each cell type as well as their structural classification determined by SQANTI. The bottom panel shows the lengths of the identified isoforms as swarmplots. Black line shows the median read length.

**Fig. 7:**
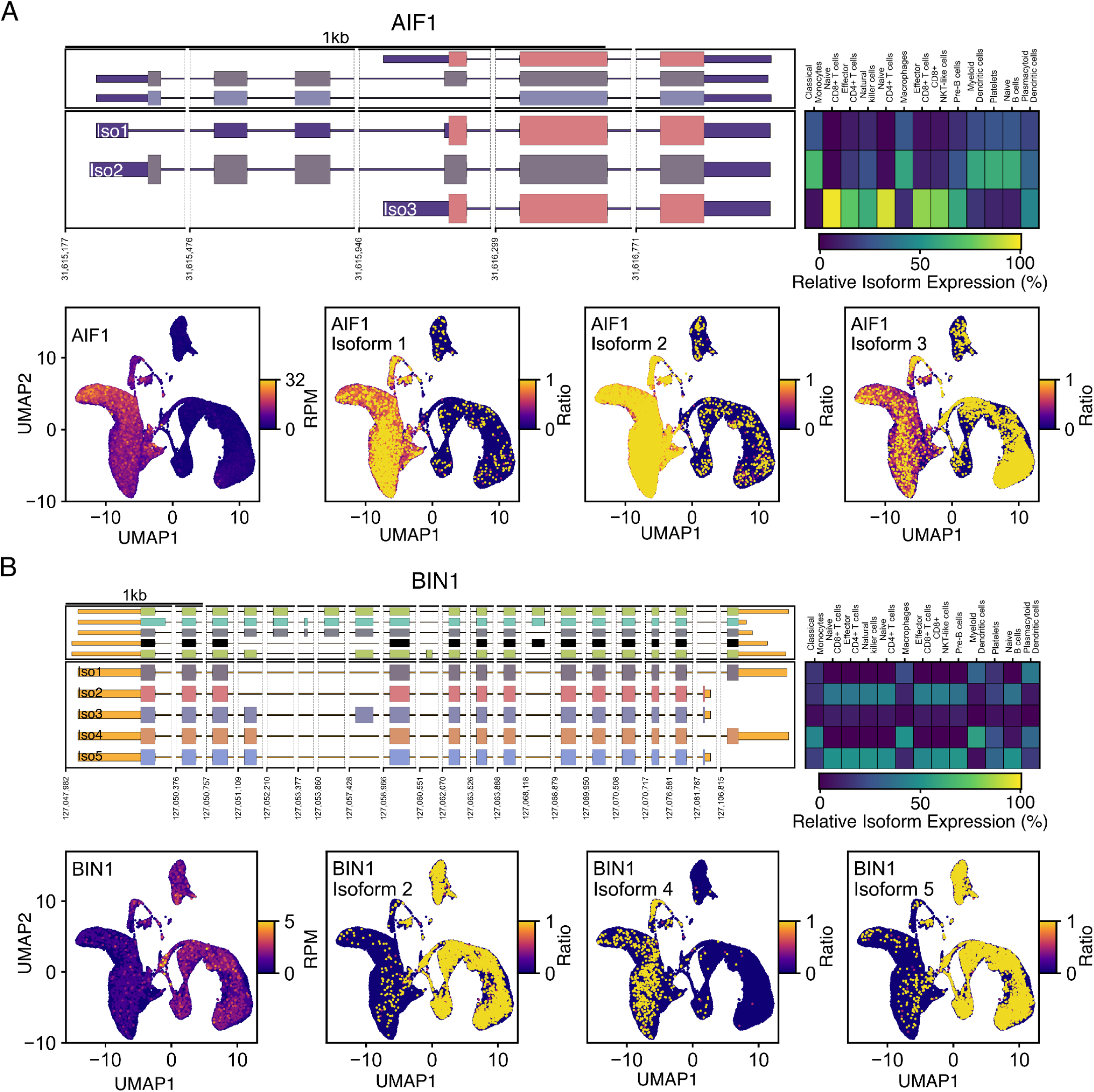
Differential isoform expression across PBMC cell types. A,B top) Genome browser representation of the AIF1 and BIN1 loci. The upper panel shows GENCODE basic annotations (subsampled for BIN1), while the lower panel shows IG-28 identified isoforms. Plus-strand features are in blue, minus-strand features in orange. Different CDSs have different random colors. x-axis shows genomic coordinates. The heatmap on the right shows the relative isoform expression of each isoform in each cell type calculated as that isoform read count divided by the sum of all isoform counts for that gene in that cell type, multiplied by 100. A, B bottom) UMAP representation of single cells from the combined sequencing runs. First panel: cells are colored based on AIF1 or BIN1 gene expression defined as the normalized total read count of AIF1 isoforms in each cell. Second, third and fourth panel: the cell’s color is determined by the relative expression of indicated isoform, defined as the isoform read count divided by the sum of read counts of all isoforms for that specific cell. Cells with high relative expression are plotted above cells with low read counts for visualization purposes.

Further analysis of the pseudo-bulk transcriptomes showed that IG-28 detected isoforms for the different cell types in a read depth dependent manner, although isoform numbers varied far less than read numbers across the different cell types (Fig 6B). Further, compared to the bulk transcriptomes in this study, the cell type pseudo-bulk transcriptomes were overall shorter (Fig. 6B, bottom) and more likely to fall within the SQANTI incomplete splice-match category (Fig. 6B, top) – both speaking to the somewhat compromised quality of fl-cDNA generated by single-cell approaches.

Because IG-28 created cell type specific bam files, we were able to provide these files to LRAA and Isocall as input and generate cell type pseudo-bulk transcriptomes with both. This allowed us to compare the pseudo-bulk transcriptomes produced by IG-28, LRAA, and Isocall – similarly to what the comparison we performed for the bulk fl-cDNA datasets (Supplementary Fig. S1). This comparison supports our previous observations: IG-28 recognises less isoforms than LRAA and Isocall and a lesser percentage of IG-28 isoforms is classified as intergenic. Around 90% of IG-28 isoforms were identified by at least one of the other tools, compared to 30% for LRAA and 40% Isocall.

This suggested that IG-28, more so than LRAA or Isocall, identified public, high-confidence isoforms when identifying pseudo-bulk transcriptomes.

### IG-28 enables differential expression analysis of isoforms across cell types in single cell data

In addition to generating pseudo-bulk transcriptomes of cell types, IG-28 also, by default, generates a combined transcriptome from samples/groups present in the dataset. In single-cell experiments, IG-28 quantifies these isoforms on the level of all single cells in the dataset. Taking advantage of this, we used a script provided with IG-28 to combine single-cell read counts into cell type read counts based on the cell-to-cell type metadata file used in module M.

We then used this cell type level quantification to identify genes that showed differential isoform expression. To do so, we first filtered genes for those with overall isoform expression of > 5 RPM in at least 2 cell types. For each gene that passed this filter, we generated a table that contained the relative isoform expression in each cell type in which the gene was expressed at > 5 RPM. Finally, we applied a chi-square test to this table. The chi-squared test produced a p-value below 0.001 (after Bonferroni multiple testing correction) for 466 of the 4314 tested genes.

This list of 466 genes included AIF1 locus, which has been known to present differential expression of isoforms in different PBMC cell types^5,29^. As in previous studies, AIF1 showed two isoform expression states roughly split between T cells and the other cell types in the dataset. Similarly, BIN1, another gene detected to feature differential isoform expression, showed two isoform expression states roughly split between monocytes, macrophages, and dendritic cells (i.e. myeloid cells) and the other cell types in the dataset.

Together, this showed that IG-28 can perform differential isoform expression analysis across cell types which will be essential to extract information from large single-cell datasets now starting to be released^30^.

## DISCUSSION

IG-28 represents an extensive rewrite of Mandalorion that minimizes RAM usage and overall processing time and enables it to identify and quantify high-confidence isoforms from large bulk and single-cell fl-cDNA datasets. We show here that it does so while matching or exceeding its predecessor’s performance.

Instead of pursuing a strategy like Isocall which processes reads from individual samples separately and merges the resulting output, IG-28 processes all reads from all samples in a dataset together – including mitochondrial genome/reads. IG-28 does so while using less than 40GB at peak, even for the 1KG dataset which contains more than 1 billion fl-cDNA reads.

This in turn means that any computer with sufficient RAM to align a long read fl-cDNA dataset using minimap2^31^ (which requires around 40GB of RAM to hold its index of the human genome) can also be used to run IG-28 on datasets of at least 1 billion reads.

IG-28 processing time increased with dataset size and complexity. Running on a desktop workstation, IG-28 took ∼20 minutes to perform the post-alignment processing WTC-11 & SIRV dataset, ∼2 hours for the Human Brain dataset, 15 hours for the 1KG dataset, and 40 hours for the scPBMC dataset (which involved calling isoforms for each cell type separately) (Fig. S2).

To evaluate the performance of IG-28 and compare it to other tools we first processed an LRGASP dataset which contained spike-in RNA transcripts. However, all tested tools performed very well in this scenario.

Therefore, for datasets that lack a ground truth, we relied on a new metric for identifying likely high-confidence isoforms (*% Public SJC)* – the percentage of isoforms (or Splice Junction Chains) a tool identifies that are also identified by any other tool. This metric is based on the fact that fl-cDNA sequencing has addressed the main shortcoming of RNA-seq. fl-cDNA reads cover transcript isoforms end-to-end which means tools should only really disagree on how they handle likely experimental artifacts and edge-cases, i.e. tools identifying isoforms from fl-cDNA data should largely agree on the isoforms they identify. In other words, most isoforms a tool identifies should be public, not private. However, LRGASP showed that this is not the case for most tools. When we calculated this metric for all tested tools and datasets in our study, we found that IG-28 had a far higher percentage of public SJCs than any other tested tool. While *% Public SJCs* reflects a tool’s ability to identify SJCs other tools have confidence in, i.e. high-confidence SJCs, it is not a perfect metric.

First, it is dependent on which group of tools are compared to each other and comparing too many tools might saturate the metric. Second, it doesn’t take into account transcript ends. While it would be straightforward to calculate public isoforms instead, thereby taking ends into account, this is complicated by the fact that tools like IsoQuant and Isocall rely heavily on genome reference annotations to determine these ends. Third, it emphasizes identifying high-confidence SJCs over identifying as many SJCs as possible which in turn might not align with the design choices of tools other than IG-28. Finally, like any other metric, it might, over time, become a target to develop towards which would undermine its utility.

After establishing its ability to identify and quantify high-confidence isoforms from bulk datasets, we showed that IG-28 also has the capability to analyze single-cell fl-cDNA datasets. We implemented this capability with the intention of being lightweight, flexible, and easy-to-use.

First, we focused on not reproducing existing solutions for tasks like cell barcode identification or UMI merging, both of which are now handled by PacBio’s software suite. Second, instead of choosing and integrating a single-cell clustering solution like either Seurat or Scanpy and providing a complex wrapper around it, IG-28 leaves this choice up to the user. IG-28 simply exports a CellxGene matrix that can be imported into the single-cell clustering workflow the user is most familiar with. The workflow can then export a cell-to-cell type table that can be imported back into IG-28. After using this cell-to-cell type table to group cells into cell types, analysis proceeds just like it would for bulk datasets, generating transcriptomes for each cell type that are quantified down to the single-cell level. Because IG-28 provides sorted bam files for each cell type, other tools can also be used to generate pseudo-bulk transcriptomes if their design choices better align with any given experiment. Further, IG-28 integrates metadata at the read name level making downstream analysis of this metadata straightforward.

In summary, IG-28 represents a tool for the analysis of fl-cDNA that excels on identifying high-confidence isoforms. IG-28 is resource-efficient and fast and should therefore be ready for the large datasets that are now starting to arrive thanks to improved sample preparation and long-read sequencing technologies.

## METHODS

### Data Generation

#### Human Brain

Human Brain RNA was purchased from Takara (Cat 636530). 200ng of total RNA was processed into fl-cDNA using a modified Smartseq2 protocol. In short, RNA was reverse transcribed using MaximaH(-) reverse transcriptase (Thermo Fisher). Reverse transcription (RT) was primed using oligodT primers (Table S1) which contained a sample index and an universal priming sequence on their 5’ end. The RT reaction also contained a template switch oligo whose sequence is integrated on the 3’ end of the resulting cDNA. The TSO sequence also contained a unique molecular identifier and a different universal priming site. The resulting fl-cDNA was amplified by PCR using primers binding the universal priming sites. The amplified cDNA was then fractionated by size using the SAGE ELF and different size fractions were processed separately using the R2C2 method.

fl-cDNA from each bin was circularized using an equal mass of DNA splint (Table S1) using NEBuilder HiFi DNA assembly master mix (NEB E2621). Splints were designed so that their ends were complementary to the ends of the cDNA. Non-circularized DNA was digested for 3 hours at 37 C using ExoI, ExoIII, and Lambda exonuclease (NEB) before heat inactivation for 20 min at 80 C. The reaction was cleaned using SPRI beads at a 0.8:1 (bead:sample) ratio. The cleaned, circularized library was then used for rolling circle amplification (RCA) using Phi 29 (NEB) with a random hexamer primer for 18 hours at 30 C before a heat inactivation for 10 min at 65 C. RCA product was debranched using T7 endonuclease (NEB) for 2 hr at 37 C and then cleaned and concentrated using a NEB Monarch DNA concentrator column. The debranched RCA product was run. The resulting R2C2 DNA was used as input into ONT library preparation.

ONT libraries were prepared from R2C2 DNA using the ONT ligation kit (SQK-LSK114) and sequenced on ONT PromethION flow cells (R10.4). Flow cells were flushed with DNAse I and loaded with additional libraries repeatedly to increase sequencing throughput.

To generate Human Brain R2C2 fl-cDNA sequences ONT raw reads for each size bin were first processed with the C3POa tools (v3.2). This generated R2C2 consensus reads. R2C2 consensus reads were then polyA trimmed with the RemovePolyA.py script of the IG-28 tool. Then trimmed R2C2 consensus reads were merged based on their UMIs using the BC1 tool (v0.95) ^32^. Finally, the UMI sequences themselves were removed from the merged, trimmed R2C2 consensus reads by discarding the first 34 nucleotides of each read.

### Data Sources and preprocessing

#### WTC-11 & SIRV

PacBio Iso-seq data was downloaded from Encode (ENCFF245IPA, ENCFF370NFS, ENCFF563QZR). Because not all files were polyA trimmed, we used the RemovePolyA.py script included in IG-28 to perform that processing step.

#### Human Brain

In addition to the R2C2 data we generated ourselves we downloaded PacBio Kinnex data from the PacBio data portal at

https://downloads.pacbcloud.com/public/dataset/Kinnex-full-length-RNA/DATA-Revio-IDT-BrainUHRR/2-FLNC/flnc-bc04-brain-regular-rep1.bam
https://downloads.pacbcloud.com/public/dataset/Kinnex-full-length-RNA/DATA-Revio-IDT-BrainUHRR/2-FLNC/flnc-bc05-brain-regular-rep2.bam
https://downloads.pacbcloud.com/public/dataset/Kinnex-full-length-RNA/DATA-Revio-IDT-BrainUHRR/2-FLNC/flnc-bc06-brain-regular-rep3.bam

#### 1KG

PacBio Kinnex fl-cDNA data of 100 lymphoblastoid cell lines was downloaded from https://s3.amazonaws.com/1000g-ont/index.html?prefix=Kinnex/kinnex_1KGP/data/processed/bam/isoseq/

This dataset is part of an ongoing 1000 Genomes long-read RNA-seq project and the full dataset is unpublished and in preparation (1000 Genomes Long-Read Sequencing Consortium).

#### scPBMCs

10X Genomics single-cell Pacbio Kinnex fl-cDNA sequences were downloaded from the following links

https://downloads.pacbcloud.com/public/dataset/Kinnex-single-cell-RNA/DATA-Revio-Kinnex-PBMC-10kcells-10xGEMX3p/2-DeduplicatedReads/scisoseq.5p--3p.tagged.refined.corrected.sorted.dedup.bam
https://downloads.pacbcloud.com/public/dataset/Kinnex-single-cell-RNA/DATA-Revio-Kinnex-PBMC-10x3p/2-DeduplicatedReads/scisoseq.5p--3p.tagged.refined.corrected.sorted.dedup.bam
https://downloads.pacbcloud.com/public/dataset/Kinnex-single-cell-RNA/DATA-Revio-Kinnex-PBMC-10x5p/2-DeduplicatedReads/scisoseq.5p--3p.tagged.refined.corrected.sorted.dedup.bam
https://downloads.pacbcloud.com/public/dataset/Kinnex-single-cell-RNA/DATA-Revio-Kinnex-PBMC-20kcells-10xGEMX3p-rep1/2-DeduplicatedReads/scisoseq.5p--3p.tagged.refined.corrected.sorted.dedup.bam
https://downloads.pacbcloud.com/public/dataset/Kinnex-single-cell-RNA/DATA-Revio-Kinnex-PBMC-20kcells-10xGEMX3p-rep2/2-DeduplicatedReads/scisoseq.5p--3p.tagged.refined.corrected.sorted.dedup.bam
https://downloads.pacbcloud.com/public/dataset/Kinnex-single-cell-RNA/DATA-Revio-Kinnex-PBMC-20kcells-10xGEMX5p/2-DeduplicatedReads/scisoseq.5p--3p.tagged.refined.corrected.sorted.dedup.bam
https://downloads.pacbcloud.com/public/dataset/Kinnex-single-cell-RNA/DATA-Vega-Kinnex-PBMC-10kcells-10xGEMX3p/2-DeduplicatedReads/scisoseq.5p--3p.tagged.refined.corrected.sorted.dedup.bam

### Isoform identification

We identified and quantified isoforms across 4 data sets (WTC-11 & SIRV, Human Brain, 1KG, scPBMCs) using Mandalorion, IG-28, FLAIR, IsoQuant, LRAA, and Isocall. Other than Mandalorion and IG-28, which require fastx or unaligned bam files, the tools required sorted bam files as input. Therefore, we used the sorted bam file generated by IG-28 as their input. Because LRAA added empty lines to its GTF file output, which broke downstream tools like SQANTI, we removed these empty lines using `grep -v ’^$’`.

#### WTC-11 & SIRV

We used the GRCh38 reference sequence and GENCODE v49 reference annotation for all tools. Additionally we added SIRV and ERCC sequences to the reference sequence and SIRV and ERCC gene models to the reference annotation. These files were downloaded from Lexogen.

Because Isocall does not accept gtfs that contain genes with isoforms on both ‘+’ and ‘-’ strands which is an edge case that occurs in the annotation of the SIRV spike-ins. We therefore modified the gene names in the provided SIRV annotation to be appended with the direction of the isoforms they contain, i.e. SIRV2 becomes SIRV2_+ and SIRV2_-.

We used the following commands to run

##### Mandalorion (v4.6)

With annotation

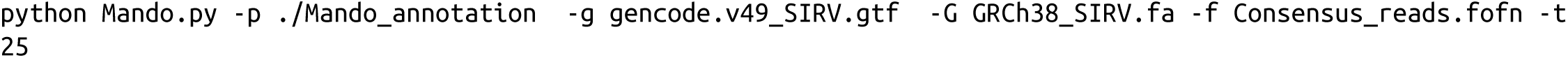

Without annotation

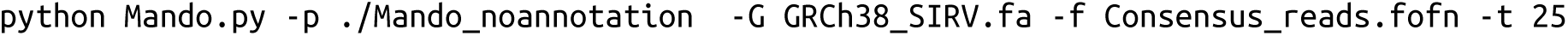

##### IG-28 (v1.0)

*With annotation*

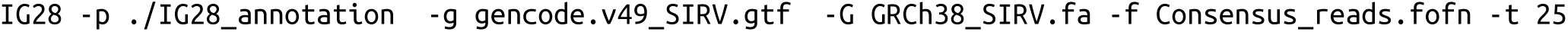

Without annotation

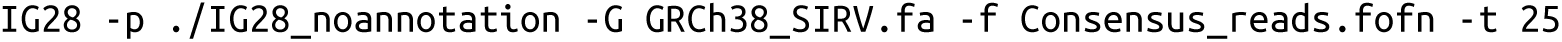

##### FLAIR (v3.0.0)

With annotation

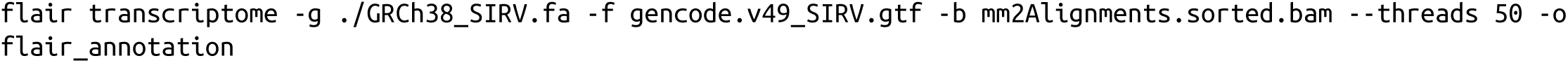

##### IsoQuant (v3.12.0)

With annotation

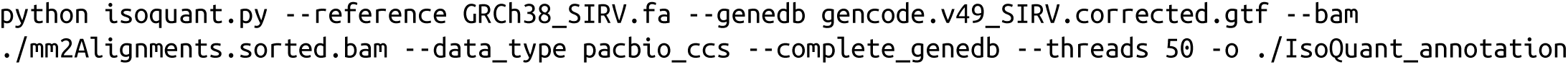

Without annotation

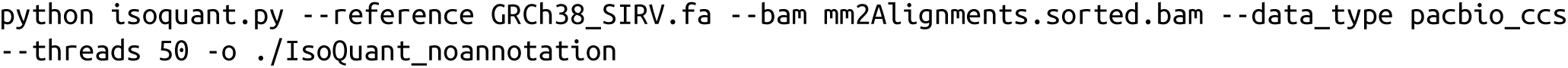

##### LRAA (v0.15.0)

With annotation

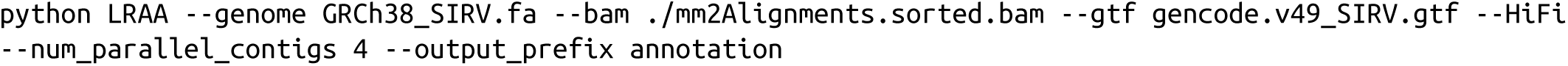

Without annotation

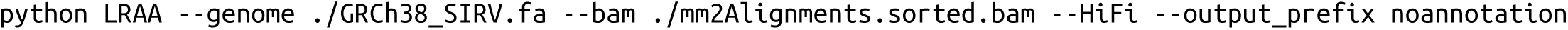

##### Isocall (v0.15.0)

With annotation

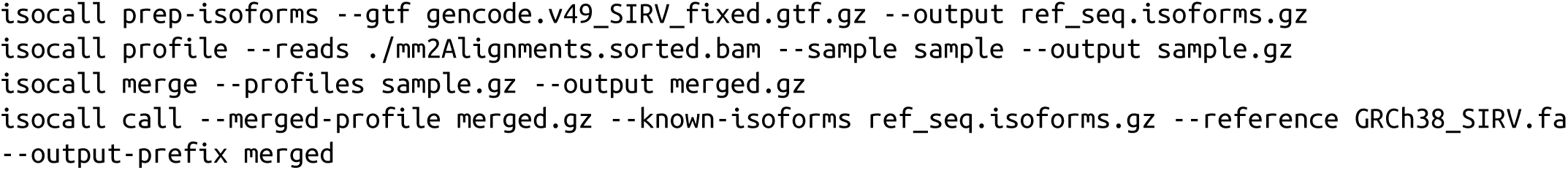

#### Human Brain

##### IG-28

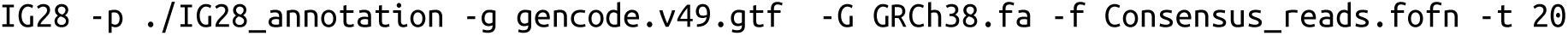

##### FLAIR

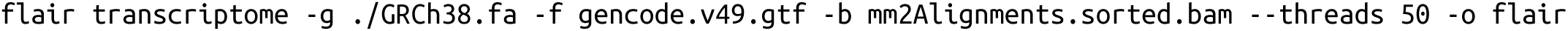

##### IsoQuant

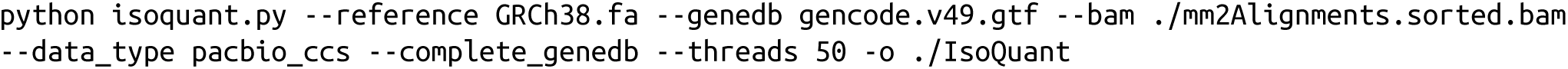

##### LRAA

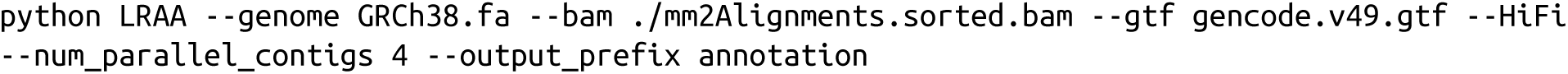

##### Isocall

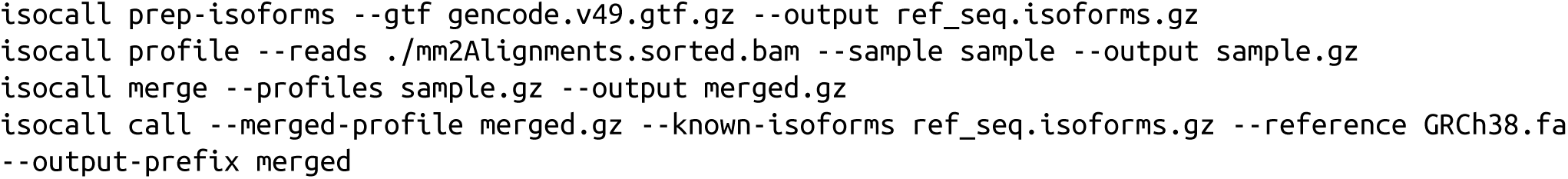

#### 1KG

##### IG-28

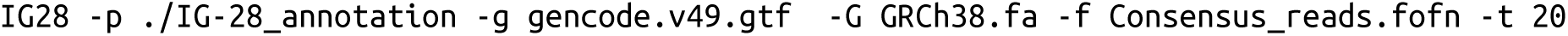

##### LRAA

Because LRAA did not complete when we attempted to run it on the entire dataset, we ran it on each chromosome separately. We let it run for ∼3 weeks until it had completed chromosome 1-10 and then terminated the run.

We used the following command structure (with chr1 as an example):

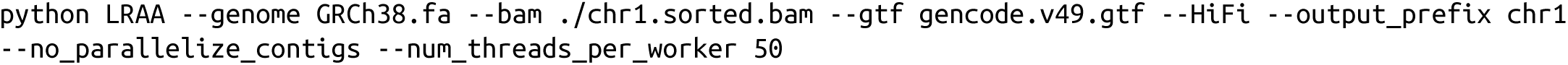

##### Isocall

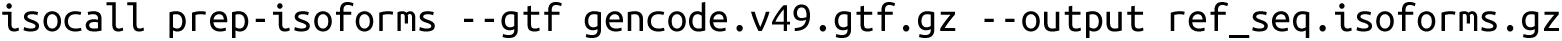

Because 100 separate samples were included in this data set which are profiled separately by Isocall by default, we first split the mm2Alignments.sorted.bam generated by IG-28 by sample into 100 separated sorted bam files named after the sample. Then, we profiled each of these files separately, like so:

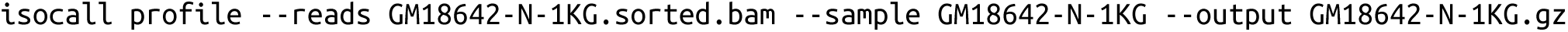

Then we merged these profiles using the following abbreviated command

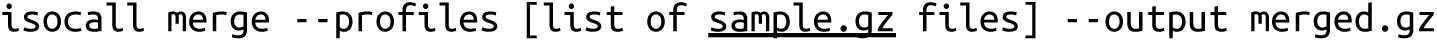

to merge all the resulting profiles and

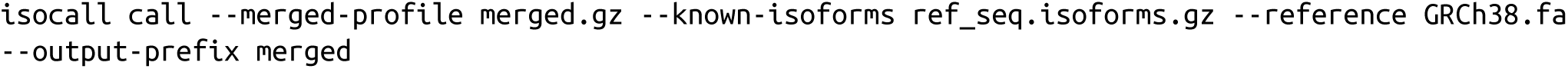

to complete the workflow.

#### scPBMCs

We first ran Module A IG-28 on the 7 scPBMCs datasets. Then, we ran module G which produce a cell x gene expression matrix. We then used that matrix to cluster and annotate single cell using Seurat. We followed the standard workflow illustrated in the Seurat single cell analysis tutorial:

https://satijalab.org/seurat/articles/pbmc3k_tutorial

We filtered cells expressing less than 350 and more than 4000 genes and with more than 20% of total counts from mitochondrial genes. We then log-normalized our data and performed batch-correction using the anchor-based CCA integration method built-in Seurat. Finally we automatically assigned cell types to each cell using the scType method. We then exported a metadata file linking each cell to its assigned cell type which was used for running module M of IG-28. Module M thus annotated the intermediate processing files and allowed IG-28 to split the full dataset into cell type-specific bam files. IG-28 then ran Module D and C to generate combined and cell type-specific isoform annotations and quantifications. Because LRAA did not complete when we attempted to run it on the entire dataset and Isocall does not have a single cell mode, we ran these tools on each cell type separately. To do so, we used the cell type specific sorted bam files produced by IG-28.

##### IG-28

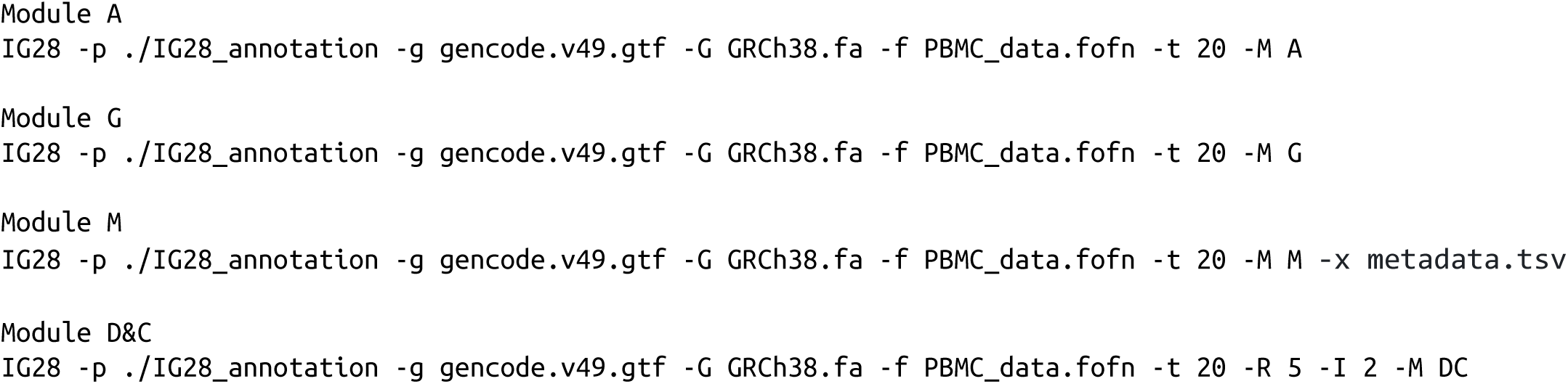

##### LRAA

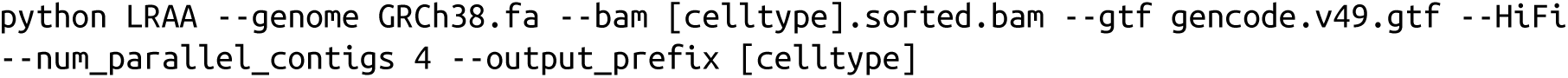

##### Isocall

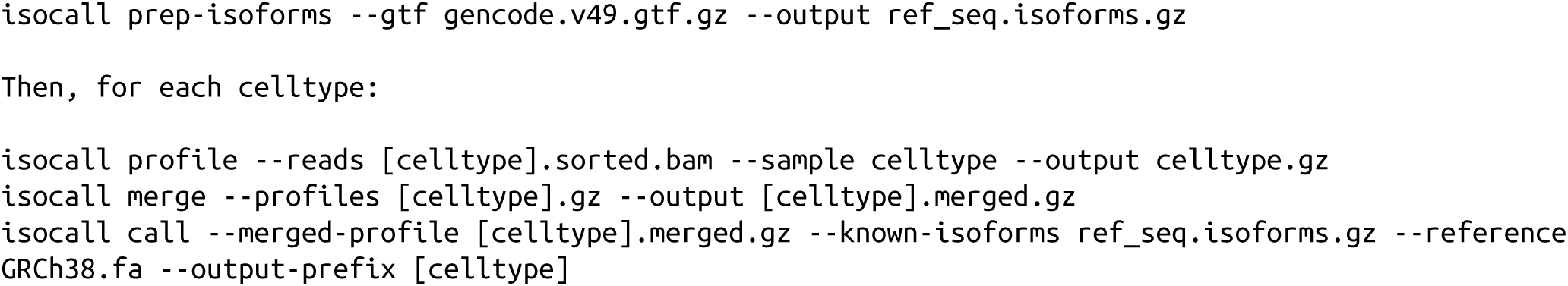

## DATA AVAILABILITY

Human Brain size-selected fl-cDNA data generated by the ONT-based R2C2 method is available at SRA under Bioproject PRJNA1497049.

## CODE AVAILABILITY

IG-28 is available at GitHub at https://github.com/christopher-vollmers/IG28.git

## FUNDING

This work has been supported by NIGMS grant R35GM133569 to C.V. and NHGRI T32HG012344 fellowship to H.E.M..

## ACKNOWLEDGEMENTS

We thank the 1000 Genomes Long-Read Sequencing Consortium, and specifically the Miller and Eichler labs at the University of Washington and the McCoy lab at Johns Hopkins University for generating and providing Kinnex full-length cDNA of 100 lymphoblastic cell lines from the 1000 Genomes Project.

## Supplement

**Figure S1:**
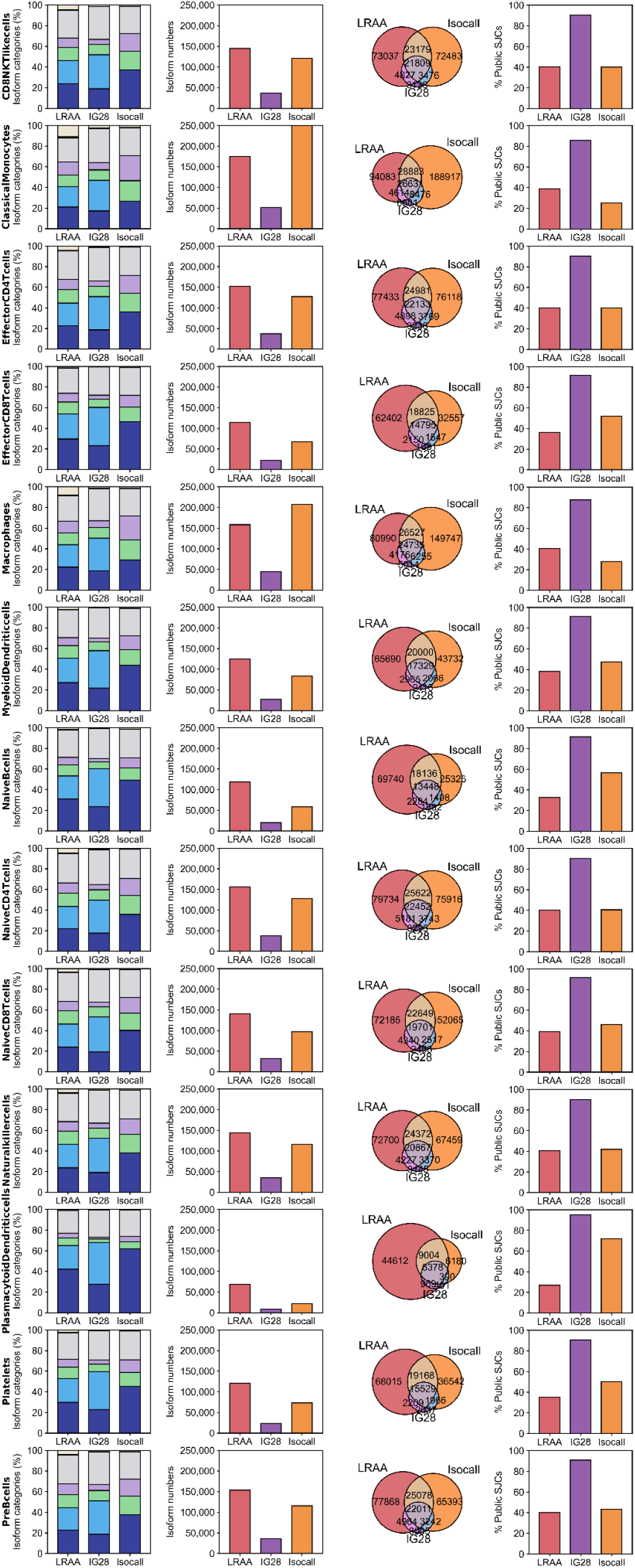
Pseudo-bulk transcriptomes called by IG-28 retain characteristics of bulk transcriptomes. Each row shows and compares the isoforms identified by IG-28, LRAA, and Isocall from the fl-cDNA Kinnex reads assigned to a distinct cell type. From left to right, the normalized structural categories, numbers, overlap of splice junction chains (SJCs), and percentage of public SJCs of the called isoforms are shown for the tree tools.

**Fig. S2:**
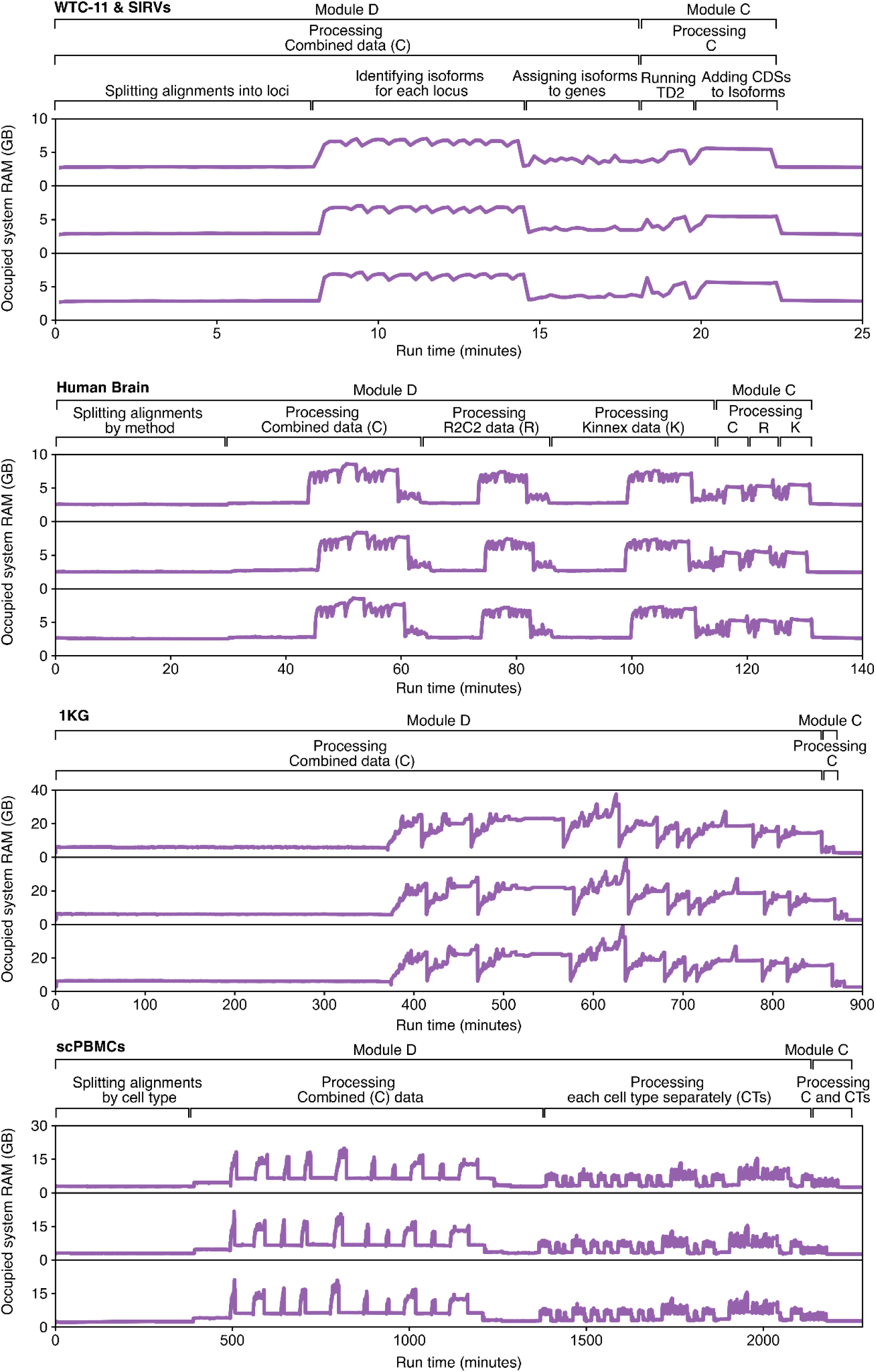
IG-28 process datasets fast and using low amounts of RAM. All 4 datasets in the study were analyzed by IG-28 in triplicate. We focused on Modules D, and C since module A contains the alignment of reads with minimap2 which is a step common to all tools and therefore part of any tools runtime.

**Table S1:**
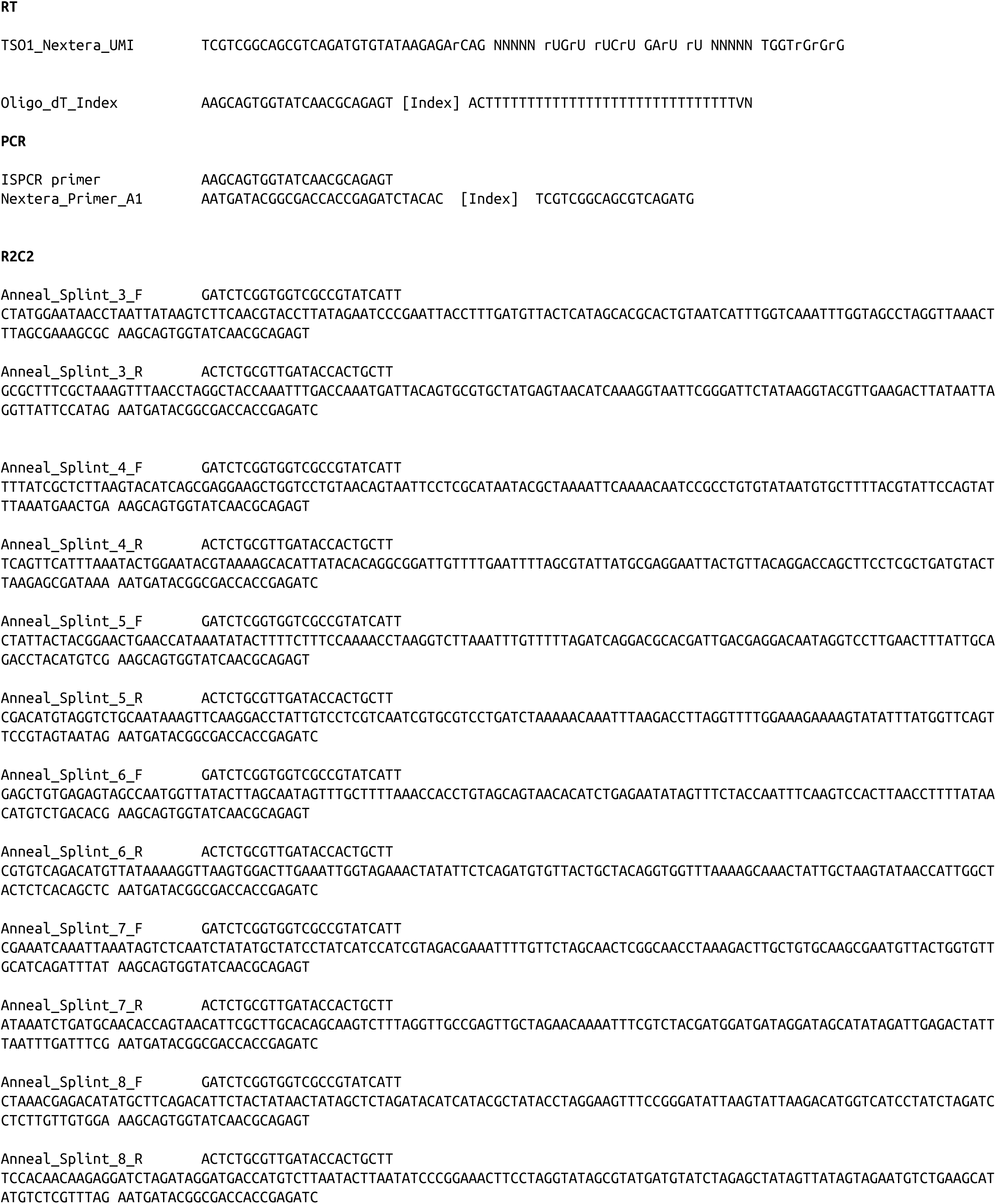
Oligos used in this study.

## Notes

### Competing Interest Statement

C.V. holds a patent on the R2C2 method used in this manuscript

